# Real-time monitoring of subcellular H_2_O_2_ distribution in *Chlamydomonas reinhardtii*

**DOI:** 10.1101/2020.11.13.382085

**Authors:** Justus Niemeyer, David Scheuring, Julian Oestreicher, Bruce Morgan, Michael Schroda

**Affiliations:** Molekulare Biotechnologie & Systembiologie, TU Kaiserslautern, Paul-Ehrlich Straße 23, D-67663 Kaiserslautern, Germany; Phytopathologie, TU Kaiserslautern, Paul-Ehrlich Straße 22, D-67663 Kaiserslautern, Germany; Institute of Biochemistry, Zentrum für Human-und Molekularbiologie (ZHMB), Saarland University, D-66123 Saarbrücken, Germany

## Abstract

H_2_O_2_ has been recognized as an important signaling molecule in plants. We sought to establish a genetically encoded, fluorescent H_2_O_2_ sensor that allows H_2_O_2_ monitoring in all major subcompartments of a Chlamydomonas cell. To this end we engineered the hypersensitive H_2_O_2_ sensor, roGFP2-Tsa2ΔC_R_, as a genetic part for the Chlamydomonas Modular Cloning toolbox. Using previously generated parts, together with new ones, we constructed modules and devices that target the sensor to the cytosol, nucleus, mitochondrial matrix, chloroplast stroma, thylakoid lumen, and ER. The sensor was functional in all compartments, except for the ER where it was fully oxidized. Employing our novel sensors, we show that H_2_O_2_ produced by photosynthetic linear electron transport (PET) in the stroma leaks into the cytosol but only reaches other subcellular compartments if produced under non-physiological conditions. Our results thus imply the establishment of steep intracellular H_2_O_2_ gradients under normal physiological conditions and suggest that the cytosolic complement of H_2_O_2_ scavenging enzymes effectively limits H_2_O_2_ diffusion. Furthermore, in heat stressed cells, we show that cytosolic H_2_O_2_ levels closely mirror temperature up- and downshifts and are independent from PET. We anticipate that these sensors will greatly facilitate future investigations into H_2_O_2_ biology in algal and plant cells.

## Introduction

In plant cells, hydrogen peroxide (H_2_O_2_), or its precursor superoxide, is produced as a side product of cellular processes including photosynthetic linear electron transport (PET), the mitochondrial respiratory chain, or substrate oxidation, for example by the photorespiration enzyme glycolate oxidase (Cheeseman, 2007; Foyer and Noctor, 2016). H_2_O_2_ is a relatively stable molecule and is largely unreactive with most proteins. It is therefore well-suited as a second messenger in cellular signaling cascades (Waszczak et al., 2018). Such signaling cascades can be fueled either with H_2_O_2_ produced as a side product, e.g. from PET (Exposito-Rodriguez et al., 2017), or with H_2_O_2_ produced specifically for a signaling cascade. One example here is the respiratory burst oxidase homolog D (RBOHD) in the plasma membrane that produces H_2_O_2_ for systemic signaling upon stimuli including wounding, cold, heat, high light and salinity (Miller et al., 2009; Lew et al., 2020), and plays a role during ABA-induced stomatal movement (Pei et al., 2000; Kwak et al., 2003).

The recognition of H_2_O_2_ as second messenger has generated a demand for tools to measure H_2_O_2_ dynamics within plant cells. The development of synthetic dyes for H_2_O_2_ detection has allowed some insights, however, several problems are inherent to their use, including variable uptake and efflux, lack of subcellular compartment specificity, lack of redox species specificity, and the irreversibility of their reaction with H_2_O_2_ (Bilan and Belousov, 2018; Roma et al., 2018). A major methodological advance in the subcellular detection of H_2_O_2_ came with the creation of genetically encoded fluorescent protein sensors that allow for subcellular compartment-specific monitoring of H_2_O_2_ in real time in living organisms. The first of these sensors was HyPer (Belousov et al., 2006). HyPer consists of a circularly permuted YFP positioned between two halves of the H_2_O_2_-sensitive *E.coli* transcription factor OxyR. The presence of H_2_O_2_ leads to the formation of a disulfide bond between the H_2_O_2_-reactive cysteine on one OxyR domain and the resolving cysteine residue on the other domain. This results in a structural change that is transmitted to the cpYFP, thereby inducing a ratiometric change in the fluorescence excitation spectrum, thus allowing measurements that are independent of probe concentration. Several improved versions of HyPer have been generated that expand its dynamic range, its oxidation and reduction rates, or employ a red fluorescent protein (Bilan and Belousov, 2018).

A second family of genetically encoded fluorescent H_2_O_2_ sensors is based on roGFP2, which contains two cysteines in adjacent β-strands on the surface of the protein β-barrel (Hanson et al., 2004). The formation of a disulfide bond between these cysteines results in small structural changes that induce a ratiometric change in the fluorescence excitation spectrum. As for HyPer, this allows measurements that are independent of sensor concentration. Since the cysteines of roGFP2 do not readily react with H_2_O_2_, roGFP2 needs to be coupled with an H_2_O_2_ reactive enzyme in a redox relay system. The first such sensor was roGFP2-Orp1, which involves a genetic fusion between the glutathione peroxidase-like enzyme Orp1 from budding yeast and roGFP2, with a short interspacing polypeptide linker (Gutscher et al., 2009). In this sensor, Orp1 sensitively reacts with H_2_O_2_ to form an intramolecular disulfide bond between its peroxidatic and resolving cysteine, followed by a thiol-disulfide exchange reaction to roGFP2.

Many cellular peroxiredoxins contain peroxidatic cysteine residues that react with H_2_O_2_ 2–3 orders of magnitude faster than the Orp1 peroxidase (Roma et al., 2018). This observation led to the generation of roGFP2-Tsa2ΔC_R_, a genetic fusion between the yeast typical 2-Cys peroxiredoxin Tsa2 and roGFP2 (Morgan et al., 2016). The resolving cysteine in Tsa2 was mutated to alanine such that reduction of the sensor by thioredoxin is impeded and sensor sensitivity further enhanced. In yeast, this sensor was shown to be approximately 20-fold more sensitive towards H_2_O_2_ than either HyPer or roGFP2-Orp1 (Morgan et al., 2016). Furthermore, roGFP2-Tsa2ΔC_R_ makes a negligible contribution to the cellular H_2_O_2_-scavenging capacity, has a low propensity for hyperoxidation, and is unaffected by changes in pH between 6 and 8.5 (Morgan et al., 2016; Roma et al., 2018). Its pH insensitivity distinguishes roGFP2-Tsa2ΔC_R_ from HyPer, for which additional pH probes need to be implemented to account for changes in pH, as observed for example in the stroma upon illumination (Schwarzlander et al., 2014; Exposito-Rodriguez et al., 2017). A pH-insensitive version of HyPer (HyPer7) has been established recently, but not yet for plant cells (Pak et al., 2020).

The aim of this work was to establish a sensitive H_2_O_2_ sensor for the unicellular green alga *Chlamydomonas reinhardtii*, a traditional model system for plant cell biology and photosynthesis (Sasso et al., 2018). To this end, we engineered roGFP2-Tsa2ΔC_R_ as a genetic part for the Chlamydomonas Modular Cloning (MoClo) toolbox for synthetic biology. We targeted the sensor to six cell compartments – cytosol, nucleus, mitochondrial matrix, chloroplast stroma, thylakoid lumen, and ER and show that changes in H_2_O_2_ concentrations can be monitored in real time in all compartments, except for the ER. Our data reveal the establishment of strong intracellular H_2_O_2_ gradients in response to physiological stresses.

## Results

### Construction of an H_2_O_2_ sensor for Chlamydomonas

To develop a genetically encoded, fluorescent H_2_O_2_ sensor for *Chlamydomonas reinhardtii*, we synthesized the coding sequence of roGFP2-Tsa2ΔC_R_ (Morgan et al., 2016) with codons optimized for expression in Chlamydomonas and containing the three *RBCS2* introns, as a level 0 part for the Chlamydomonas Modular Cloning (MoClo) kit (Crozet et al., 2018) (Figure 1A). This part was then assembled into level 1 modules which included either the *PSAD* promoter or the *HSP70A-RBCS2* fusion promoter (pAR), an *RPL23* terminator, and various N-terminal targeting peptides, C-terminal tags, or retention signals to facilitate localization to defined subcellular compartments. The resulting level 1 modules were then assembled into level 2 devices together with a spectinomycin resistance (*aadA*) cassette controlled by the *PSAD* promoter and terminator (Figure 1B). The level 2 devices were then transformed into the UVM4 expression strain (Neupert et al., 2009). As a control, we generated a level 0 part containing only the roGFP2 coding sequence, harboring the first *RBCS2* intron, and placed it under control of the *AR* promoter (roGFP2, Figure 1B). The best promoters and targeting peptides to use in Chlamydomonas is still a subject of debate. We therefore tested different promoters and targeting peptide combinations. For each device, at least 13 spectinomycin resistant transformants were picked and screened for sensor expression by immunoblotting. Between 30% and 70% of the transformants expressed the sensor to readily detectable levels (Supplemental Figure 1). To compare the expression levels of the sensors, whole-cell proteins of the best-expressing transformants (Supplemental Figure 1) were analyzed on the same immunoblot using an antibody against GFP (Figure 1C).

**Figure 1.**
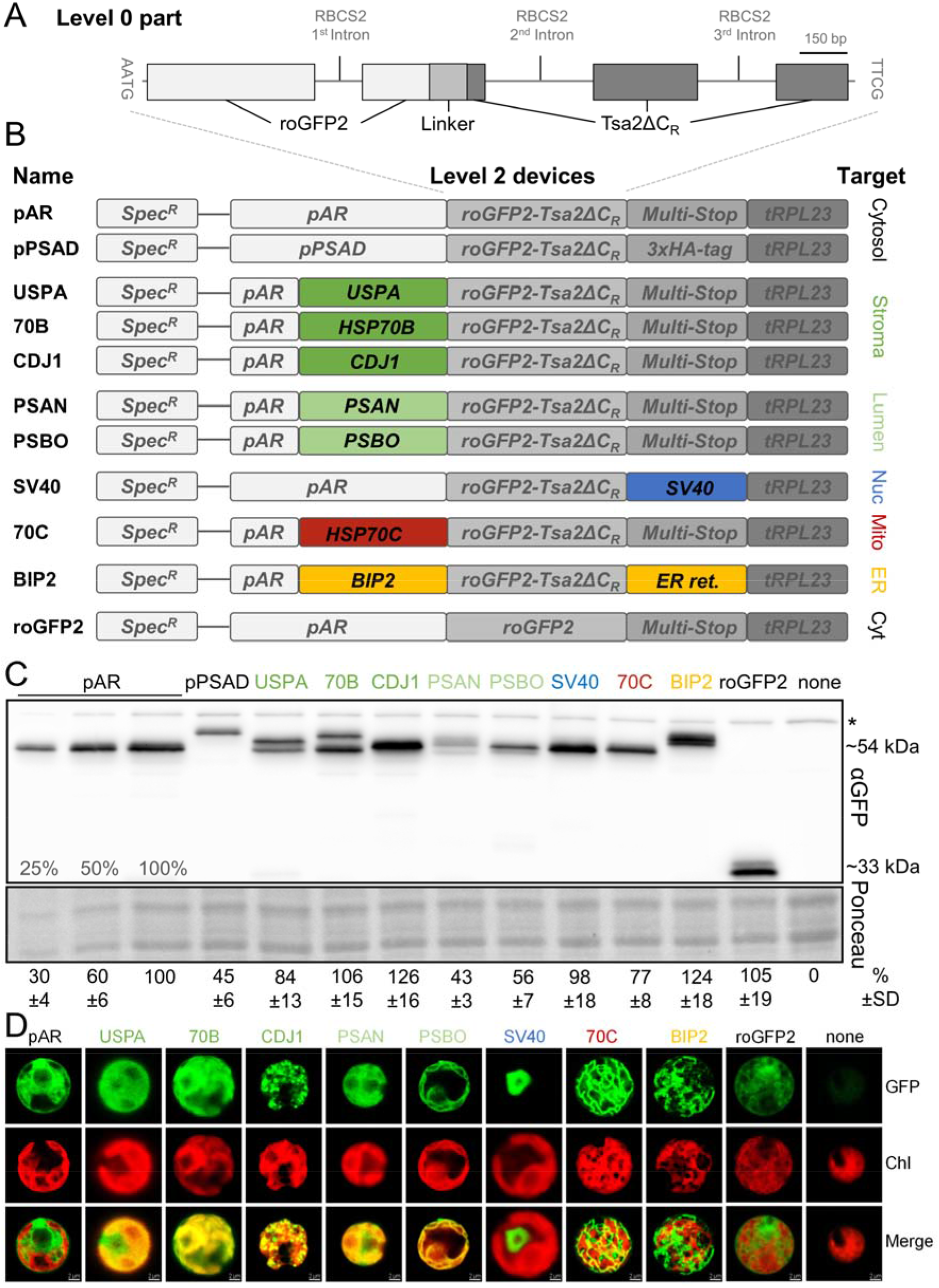
Targeting roGFP2 sensors to various compartments of a Chlamydomonas cell. **(A)** Level 0 part of the roGFP2-Tsa2ΔC_R_ H_2_O_2_ sensor. The 2132-bp ORF (exons shown as boxes), interrupted by the three RBCS2 introns (thin lines), was synthesized with optimal Chlamydomonas codon usage. RoGFP2 (light grey) is separated from Tsa2ΔC_R_ (dark grey) by a Linker (medium grey). **(B)** Using MoClo, the roGFP2-Tsa2ΔC_R_ part or only the roGFP2 part were equipped with the *HSP70A-RBCS2* promotor (*pAR*) or the *PSAD* promoter (*pPSAD*) and the *RPL23* terminator (*tRPL23*) as well as various N- and C-terminal targeting sequences to target the sensors to the cytosol (cyt), the stroma, the thylakoid lumen (lumen), the nucleus (nuc), mitochondria (mito), or the endoplasmic reticulum (ER). The resulting level 1 modules were combined with another level 1 module comprising the *aadA* resistance marker flanked by the *PSAD* promoter and terminator (Spec^R^) to yield the eleven level 2 devices shown. (C) Comparison of roGFP2 expression levels in transformants generated with the level 2 devices shown in (B). The transformants shown represent the ones with highest expression levels among at least 13 transformants screened per construct. Total cell proteins corresponding to 1.5 μg chlorophyll of each transformant and the UVM4 recipient strain (none) were separated by SDS-PAGE and analyzed by immunoblotting using an antibody against GFP. Signals retrieved from three independent experiments were quantified and normalized to the signal obtained with the transformant harboring the construct for the expression of cytosolic roGFP2-Tsa2ΔC_R_ driven by the *HSP70A-RBCS2* promotor (pAR), which was set to 100%. Mean values are given below the panel (± SD). A representative experiment is shown, with Ponceau staining demonstrating equal loading. The asterisk indicates a non-specific cross-reaction with the GFP antibody. (D) Confocal microscopy images of individual cells of the best-expressing transformants shown in (C) and the untransformed UVM4 strain (none). Shown are GFP fluorescence, chlorophyll fluorescence (Chl) and both signals merged.

With respect to cytosolic-localized constructs, we found that expression of the sensor driven by the pAR promoter, with omission of the C-terminal 3xHA tag, was ~2-fold higher than the *PSAD* promoter-driven construct (Figure 1C). Furthermore, unfused roGFP2 was expressed at about the same level as the full sensor (Figure 1C). For targeting to the mitochondrial matrix, ER, and nucleus we employed HSP70C (heat shock protein 70C) and BiP2 targeting sequences as well as an SV40 (Simian Virus 40) nuclear localization signal. For each construct, we found single protein bands in immunoblots and correct sensor localization as confirmed by confocal fluorescence microscopy (Figures 1C and 1D). For targeting to the chloroplast stroma, we tested three different transit peptides derived from USPA (universal stress protein A), HSP70B, and CDJ1 (chloroplast DnaJ homolog). A double band was observed in all USPA transformants and some GFP fluorescence was observed in the cytosol (Figures 1C and 1D; Supplemental Figure 1). We therefore reasoned that the USPA transit peptide was not efficiently driving chloroplast import. Likewise, as judged from the protein double band and residual GFP fluorescence in the cytosol, the HSP70B transit peptide was also unable to efficiently drive sensor import into chloroplasts (Figures 1C and 1D). In contrast, using the CDJ1 transit peptide, we observed a single protein band in Western blots and an exclusive chloroplast localization judged by fluorescence microscopy (Figures 1C and 1D). We also wanted to target the sensor to the thylakoid lumen and for this created level 0 parts encoding the bipartite signaling sequences from the PSAN and PSBO proteins, which are nuclear-encoded subunits of photosystem I and II, respectively. PSAN uses the TAT pathway for the import of folded proteins, while PSBO uses the Sec pathway for the import of unfolded proteins (Albiniak et al., 2012). Immunological detection of the sensor in the best-expressing PSAN and PSBO transformants showed that the sensor accumulated only to about half of the levels detected in the pAR transformant. Moreover, in PSAN transformants, multiple protein bands were detected that migrated with larger apparent mass than expected for the mature protein (Figure 1C, Supplemental Figure 1). GFP fluorescence was detectable all over the chloroplast and even in the cytosol (Figure 1D), indicating incomplete import of the sensor into both, chloroplast and thylakoid lumen by the PSAN bipartite targeting sequence. In contrast, in PSBO transformants only one major protein band migrating with the expected mass was detected and GFP fluorescence was restricted to the thylakoid lumen, indicating full functionality of the PSBO bipartite targeting signal (Figures 1C and 1D, Supplemental Figure 1).

### RoGFP2-Tsa2ΔC_R_ functions as an ultra-sensitive H_2_O_2_ sensor in Chlamydomonas

We first tested whether we could detect the cytosolic roGFP2-Tsa2ΔC_R_ sensor and monitor its oxidation by exogenously added H_2_O_2_ in real time. To this end, we took cells of the best-expressing pPSAD transformant. Cells were concentrated by centrifugation in a 96 well microtiter plate and GFP fluorescence was monitored after adding 0, 0.1, 0.5, and 1 mM H_2_O_2_ in a fluorescence plate reader. Note that these H_2_O_2_ concentrations are initial exogenous concentrations; in HeLa cells, the resulting intracellular H_2_O_2_ concentrations have been estimated to be several hundred-fold lower (Huang and Sikes, 2014). As shown in Figure 2A, we could indeed detect sensor oxidation in a dose-dependent manner, but the signal was very noisy. In contrast, a clear H_2_O_2_ concentration-dependent sensor response was observed in the pAR transformant (Figure 2B), presumably because sensor expression was more than two-fold higher than in the pPSAD transformant. The cytosolic roGFP2-Tsa2ΔC_R_ sensor was ~20% oxidized at steady state at the beginning of our assays. Upon H_2_O_2_ addition, we observed a rapid probe oxidation, which slowly recovered over a period of ~80 min, presumably in a GSH/glutaredoxin-dependent manner as previously shown in yeast (Morgan et al, 2016). The unfused cytosolic roGFP2 was completely unresponsive to H_2_O_2_ (Figure 2C), confirming the requirement of the Tsa2 moiety to couple roGFP2 oxidation with changes in H_2_O_2_ availability. Interestingly, the kinetics of roGFP2-Tsa2ΔC_R_ oxidation and reduction were very similar to those observed in yeast (Morgan et al., 2016). This included a reduction of sensor oxidation over time in the untreated samples, which in yeast was shown to be caused by cell-dependent oxygen consumption from the assay buffer. This observation strongly suggests that roGFP2-Tsa2ΔC_R_ functions as an ultra-sensitive sensor in Chlamydomonas and is capable of sensing and responding to changes in basal cytosolic H_2_O_2_ levels, as it does in yeast.

**Figure 2.**
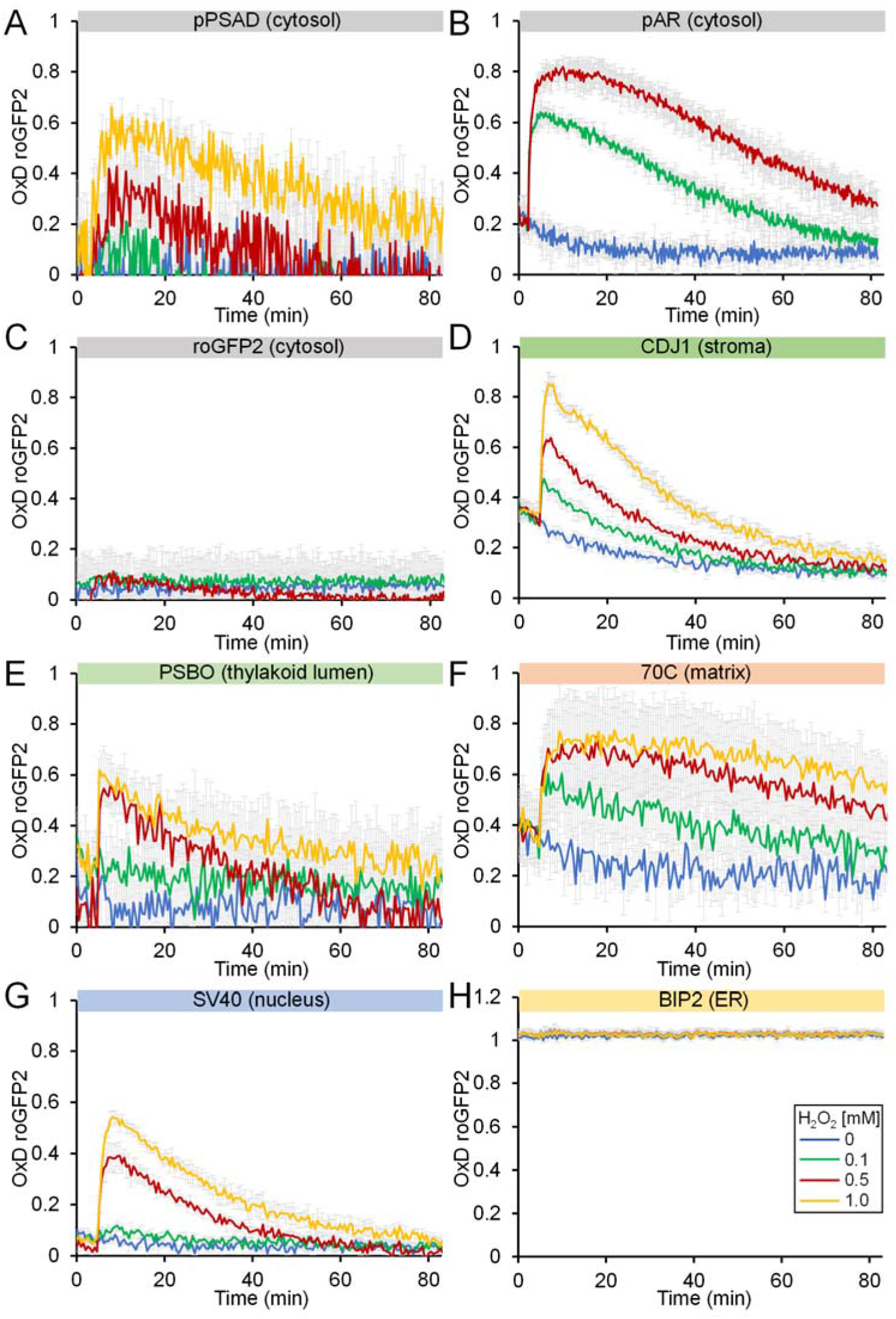
Real-time monitoring of H_2_O_2_ levels in different subcellular compartments under steady-state conditions and after the addition of H_2_O_2_. **(A-H)** Fluorescence measurement of roGFP2-Tsa2ΔC_R_ and roGFP2 in transformant cells shown in Figure 1C and 1D under steady-state conditions (no H_2_O_2_ added, blue) and after the addition of H_2_O_2_ at concentrations of 0.1 mM (green), 0.5 mM (red), and 1 mM (yellow). Values were calculated relative to those obtained for fully reduced (0) and fully oxidized (1) sensors. Shown are means of three independent experiments, error bars represent standard deviations.

Having confirmed the feasibility of measurements with cytosolic roGFP2-Tsa2ΔC_R_, we next tested the response to exogenously added H_2_O_2_ of the sensors targeted to the five additional cellular compartments. Under steady state conditions, stromal and mitochondrial sensors were more oxidized than those in the cytosol and nucleus (Figures 2B, D, F, G). However, as for the cytosolic sensor, in response to exogenous H_2_O_2_ we observed a rapid dose-dependent oxidation followed by slow reduction, for sensors targeted to the chloroplast stroma, thylakoid lumen, mitochondrial matrix and the nucleus (Figure 2D–G). Measurements were not possible in the ER as the sensor was found to be fully oxidized at steady state (Figure 2H). Signal noise inversely correlated with the expression level of the sensor: the highest noise was detected for the thylakoid lumen sensor, which was expressed at about two-fold lower levels than the sensors in the stroma and nucleus, which showed least noise (Figures 1C and 2D, E, G). The nuclear, stromal, and thylakoid lumen-localized sensors were reduced at faster rates than their counterparts in the cytosol and mitochondrial matrix (Figures 2B, D-G). Overall, the cytosolic sensor exhibited the highest sensitivity to exogenously added H_2_O_2_ and the nuclear sensor the lowest, as judged from their responsiveness to the lowest H_2_O_2_ concentration employed (0.1 mM) (Figure 2B, G). This result may indicate that H_2_O_2_ scavenging enzymes in the cytosol limit the diffusion of exogenous H_2_O_2_ to the intracellular organelles

### RoGFP2-Tsa2ΔC_R_ reveals the formation of light-dependent subcellular H_2_O_2_ gradients

Having established Chlamydomonas lines with sensors in different subcellular compartments, we asked whether we could detect changes in H_2_O_2_ levels under challenging environmental conditions. We first tested whether exposing cells to high light resulted in increased H_2_O_2_ levels in the chloroplast stroma and whether H_2_O_2_ could diffuse into other subcellular compartments. The fluorescence assay used so far allowed monitoring the sensor oxidation state in real time. However, as we had no way to illuminate cells in our plate reader assay, we turned to a ‘redox trapping’ approach. We adopted a protocol from yeast that allows for a rapid ‘trapping’ of the sensor oxidation state with the membrane permeable alkylating agent N-ethylmaleimide (NEM) (Morgan et al., 2016). NEM irreversibly alkylates free thiol groups, thus preventing further probe oxidation or reduction. We grew transformants expressing the sensor in the cytosol, nucleus, matrix, stroma, and thylakoid lumen under low light of 30 μmol photons m^−2^ s^−1^ to mid-log phase. Cultures were then kept at low light or exposed to 1000 μmol photons m^−2^ s^−1^ in the absence or presence of 3- (3,4-dichlorophenyl)-1,1-dimethylurea (DCMU) to block PET. Cells were harvested before and during the time course of the treatment and the oxidation state of the sensor was measured by plate reader after NEM-trapping. Upon shifting cells into high light, H_2_O_2_ levels in the stroma increased rapidly within minutes and remained at higher levels when compared with cells kept under low-light during the 20-min time course (Figure 3A). The cytosolic probe also responded rapidly, although probe oxidation appeared to recover after 20 min. With slower kinetics, we also observed a slight, non-significant trend towards higher H_2_O_2_ levels in high light in the mitochondrial matrix, while no increase could be detected in the nucleus (Figures 3C and 3D). The increase in H_2_O_2_ levels in all tested compartments was abolished when cells were exposed to high light in the presence of DCMU, indicating that H_2_O_2_ in high light was produced by PET. Because of its low signal to noise ratio (Figure 2E), we were unable to monitor the response of the thylakoid lumen sensor in this assay.

**Figure 3.**
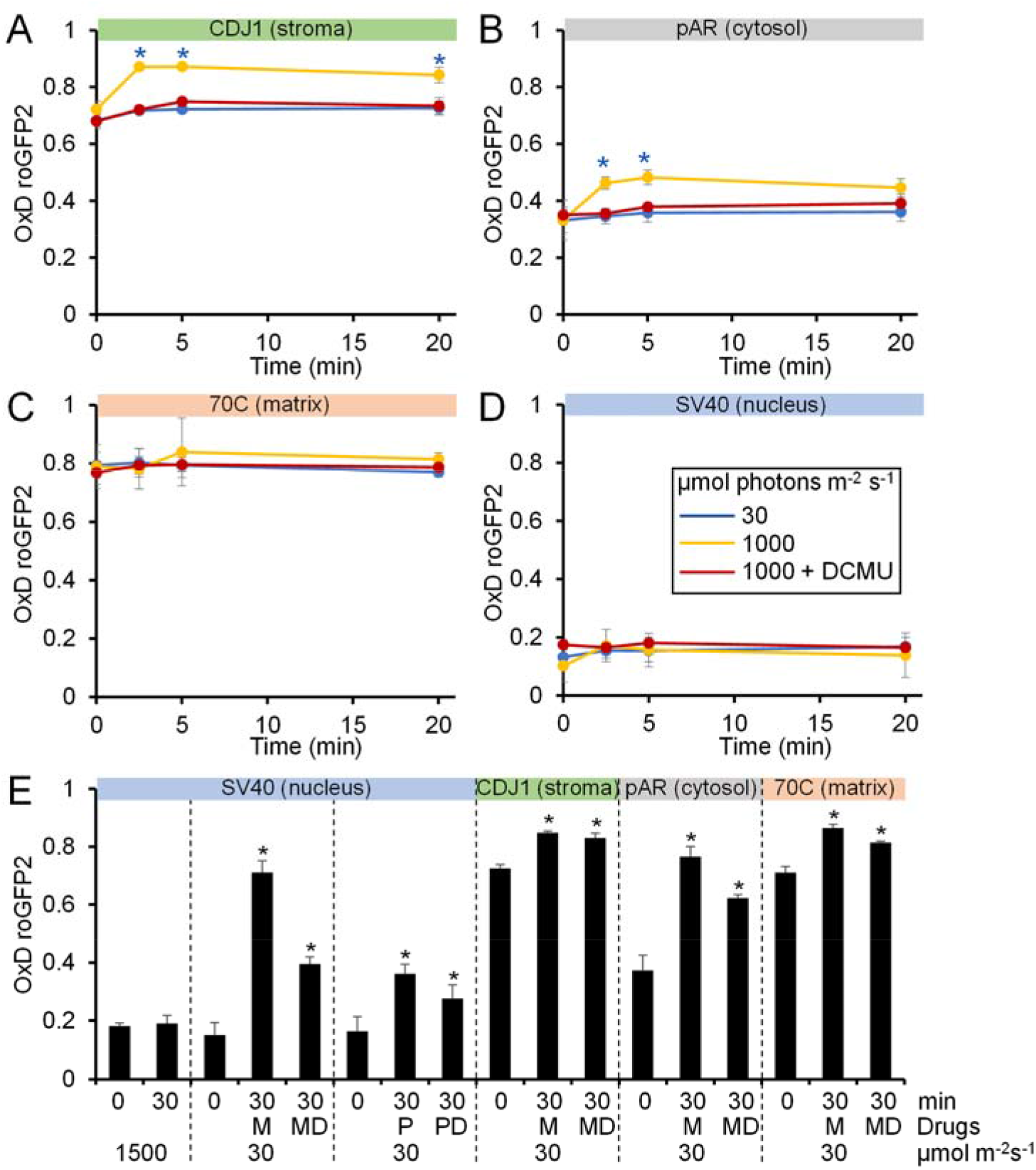
Monitoring H_2_O_2_ levels in different subcellular compartments under high-light exposure and the application of metronidazole and paraquat. Chlamydomonas transformant cells, expressing roGFP2-Tsa2ΔC_R_ in the stroma **(A)**, cytosol **(B)**, mitochondrial matrix **(C)**, and nucleus **(D)** were kept in low light of 30 μmol photons m^−2^ s^−1^ (blue) or exposed to high light of 1000 μmol photons m^−2^ s^−1^ in the absence (yellow) or presence (red) of 10 μM DCMU. The oxidation state of the sensor was trapped by the addition of NEM and roGFP2 fluorescence was measured in a plate reader. Shown are mean values from three independent experiments, error bars represent standard deviations. Asterisks indicate significant differences with respect to the low light control (t-Test, p < 0.01). **(E)** Transformant cells expressing roGFP2-Tsa2ΔC_R_ in the compartments indicated were grown in low light of 30 μmol photons m^−2^ s^−1^ (0) and exposed to 1500 m^−2^ s^−1^ for 30 min, or kept at low light for 30 min in the presence of 2 mM metronidazol (M) or 1 μM paraquat (P), both in the absence or presence of 10 μM DCMU (D). The oxidation state of the sensor was trapped by the addition of NEM and roGFP2 fluorescence was measured in a plate reader. Error bars represent standard deviations from three independent experiments. Asterisks indicate significant differences with respect to the low light control (t-Test, p < 0.05).

That we could not detect any increase of H_2_O_2_ levels in the nucleus of Chlamydomonas cells exposed to high light appeared surprising because such an increase was previously observed in *Nicotiana benthamiana* epidermal cells exposed to 1000 μmol photons m^−2^ s^−1^ (Exposito-Rodriguez et al., 2017). We therefore repeated our experiment at higher light intensities of 1500 μmol photons m^−2^ s^−1^ for 30 min but were still not able to detect an increase of H_2_O_2_ levels in the nucleus (Figure 3E).

Metronidazole and paraquat, also known as methyl viologen, both facilitate the Mehler reaction in intact cells, i.e., the transfer of electrons from PSI to oxygen to produce superoxide, which is converted to H_2_O_2_ by superoxide dismutase (Mehler, 1951; Asada, 2000). While paraquat is directly reduced by PSI, metronidazole is reduced by ferredoxin (Schmidt et al., 1977). Paraquat is active at much lower concentrations than metronidazole. We employed both drugs to test, whether the enhanced rates of H_2_O_2_ produced in their presence even under low light intensities can be detected in the nucleus. As shown in Figure 3E, this was indeed the case. We also monitored H_2_O_2_ production upon metronidazole treatment in the stroma, cytosol, and matrix and found H_2_O_2_ levels to increase in all these subcompartments. In all cases, DCMU reduced the accumulation of H_2_O_2_ but did not abolish it, most likely because plastoquinone reduction here occurs from starch breakdown via the Ndh2 protein mediating so-called pseudo-linear electron transport (Bulté et al., 1990; Mus et al., 2005; Desplats et al., 2009). This was also observed in Arabidopsis seedlings treated with paraquat and DCMU in the light (Ugalde et al., 2020).

In summary, our data show a rapid, PET-dependent increase in H_2_O_2_ levels in the chloroplast stroma. While H_2_O_2_ levels produced by metronidazole or paraquat feeding were so high that H_2_O_2_ readily diffused into all other subcompartments, H_2_O_2_ production under high light was of sufficient magnitude to increase cytosolic H_2_O_2_ but did not impact significantly upon H_2_O_2_ levels in the mitochondrial matrix or nucleus. Thus, these results are consistent with our exogenous H_2_O_2_ application experiments, suggesting that the cytosol acts as an effective barrier, which limits the intracellular diffusion of H_2_O_2_ concentrations typically produced under physiological conditions.

### Heat stress derived H_2_O_2_ is independent of photosynthetic electron transport

We chose heat stress as a second environmental challenge. To test whether H_2_O_2_ levels within Chlamydomonas cells changed under heat stress, we employed the cell line expressing the H_2_O_2_ sensor in the cytosol, as this sensor exhibited the highest signal-to-noise ratio (Figure 2B). To allow us to monitor real-time changes in cytosolic H_2_O_2_ during heat stress, we developed an experimental setup in which cells were cultivated under agitation and continuously circulated to a quartz cuvette inserted into a spectrofluorometer via connected tubing and a peristaltic pump. The culture flask, equipped with a temperature sensor, was kept on a platform well above the water surface of a 40°C water bath, placed into the water bath 20 min after the fluorescence recording had started, and placed back onto the platform another 30 min later for recovery. Interestingly, sensor oxidation and thus H_2_O_2_ production occurred immediately after the temperature had increased and immediately declined when the temperature had dropped again (Figure 4A). When the initial starting temperature was reached after 30 min of recovery, probe oxidation had not fully recovered, yet. To verify that the change in fluorescent light emission under heat stress was not caused by a heat-induced conformational change of the sensor, we trapped the sensor with NEM in steady state as well as in the fully oxidized and fully reduced states and recorded fluorescent light emission after excitation at 405 and 488 nm at 25°C, 30°C, 35°C, 40°C, and 45°C. As shown in Supplemental Figure 2, we observed no change in fluorescence emission intensities, indicating that in this temperature range the sensor is stable and that the changes in fluorescence emission under heat stress are indeed caused by changes in roGFP2 oxidation as a result of changes in cellular H_2_O_2_ levels.

**Figure 4.**
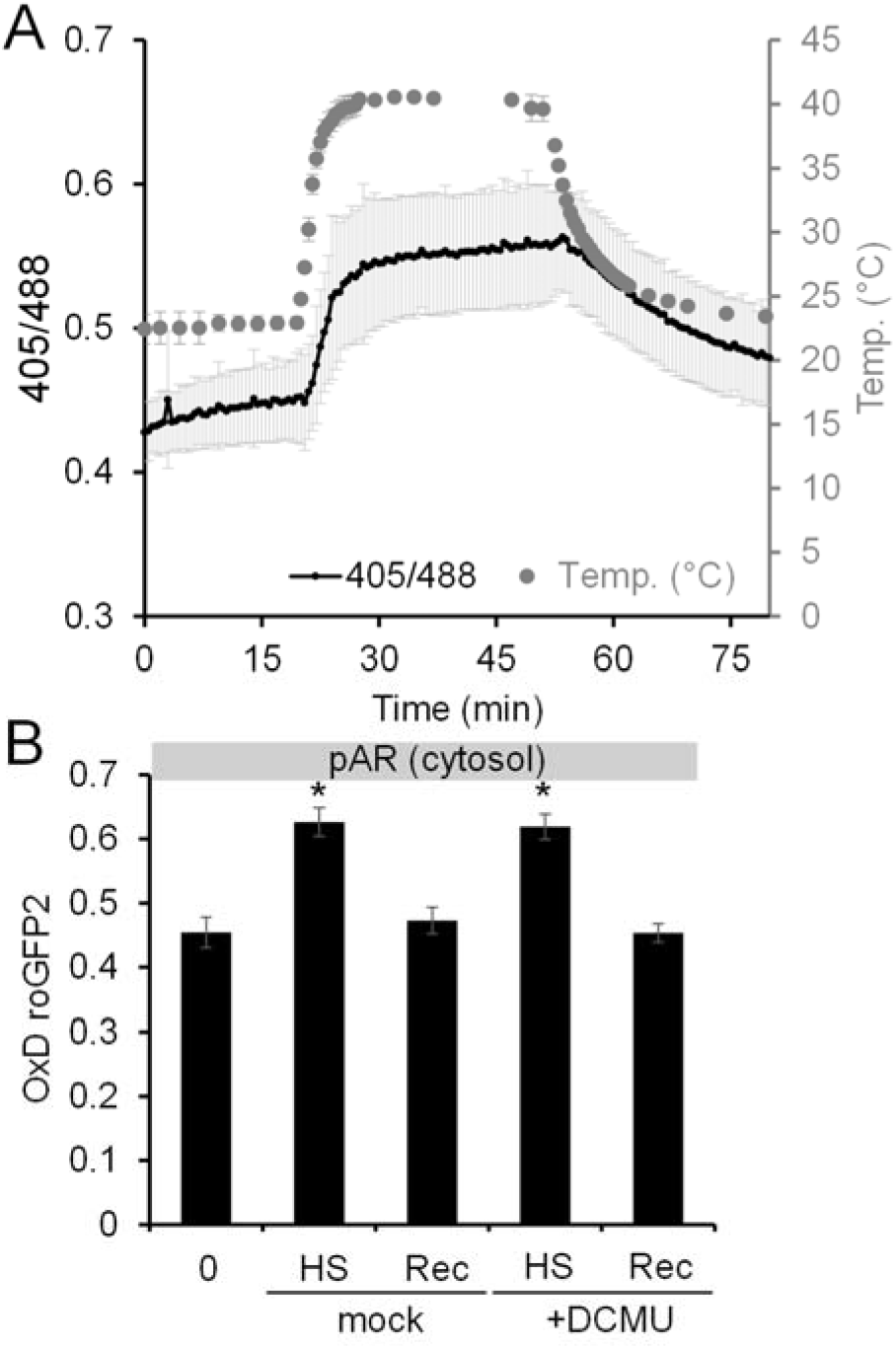
Monitoring H_2_O_2_ levels in the cytosol during heat stress and recovery. **(A)** Chlamydomonas cells expressing roGFP2-Tsa2ΔC_R_ in the cytosol were grown in low light of 30 μmol photons m^−2^ s^−1^ for 20 min, exposed to 40°C for 30 min, and shifted back to 23°C for 30 min. The ratio of the fluorescence light emitted after excitation at 405 and 488 nm, respectively, was monitored in real time using a spectrofluorometer (black line). The temperature in the culture was monitored in parallel (grey circles). Shown are mean values from three independent experiments, error bars represent standard deviations **(B)** Transformant cells expressing roGFP2-Tsa2ΔC_R_ in the cytosol were grown in low light of 30 μmol photons m^−2^ s^−1^ (0) at 23°C, exposed to 40°C for 30 min (HS), and shifted back to 23 for another 30 min (Rec). Cultures were supplied with 10 μM DCMU or only the solvent (mock) before the experiment was started. The oxidation state of the sensor was trapped by the addition of NEM and roGFP2 fluorescence was measured in a plate reader. Error bars represent standard deviations from three independent experiments. Asterisks indicate significant differences with respect to the 23°C control (t-Test, p < 0.001).

As a further test of our observations, we used the same setup employed for the high light experiments (Figure 2). Specifically, samples were taken from a culture expressing the cytosolic sensor before shifting the temperature from 23°C to 40°C, when the culture was exposed to 40°C for 30 min, and 30 min after shifting the culture back to 23°C. The redox state of the sensor was measured after trapping with NEM. As shown in Figure 4B, we could confirm that heat stress leads to a transient increase in cytosolic H_2_O_2_ levels, which was abolished when the temperature dropped again. As the increase in H_2_O_2_ levels was the same in the presence of DCMU (Figure 4B), we can conclude that H_2_O_2_ produced during heat stress does not derive from PET.

## Discussion

### RoGFP2-Tsa2ΔC_R_ is a novel sensor for the detection of H_2_O_2_ in five Chlamydomonas compartments

Here we report on the development of Chlamydomonas lines expressing the ultra-sensitive, genetically encoded H_2_O_2_ sensor roGFP2-Tsa2ΔC_R_ (Morgan et al., 2016) in five different subcellular compartments (cytosol, nucleus mitochondrial matrix, chloroplast stroma, and thylakoid lumen). The establishment of the sensor in Chlamydomonas was facilitated by using the Modular Cloning (MoClo) technology (Weber et al., 2011) and the recently generated Chlamydomonas MoClo kit (Crozet et al., 2018). The interchangeability of individual parts and the ability to assemble them in a single reaction, combined with the short generation time of Chlamydomonas, enabled iterative cycles of construct production and testing in a very short time frame. This way, problems encountered with insufficient promoter activity or inefficient targeting to the chloroplast and thylakoid lumen could be rapidly solved.

We demonstrate the suitability of roGFP2-Tsa2ΔC_R_ to monitor intracellular H_2_O_2_ levels in Chlamydomonas with three different experimental setups, each having inherent advantages and disadvantages. First, we developed an assay using microtiter plates in a plate reader (Figure 2). The advantage here is that changes in H_2_O_2_ levels can be monitored in real-time and with high throughput. The disadvantage of this assay is that cells need to be concentrated by centrifugation and we have no way to use this approach for the monitoring of responses to changes in light intensity. Second, we continuously circulated cells between culture flask and a cuvette in a spectrofluorometer (Figure 4A). The advantage of this setup is that changes in H_2_O_2_ levels can be monitored in real-time under physiological conditions (if the dark period in the cuvette is not a problem). The disadvantage is the low throughput. Third, we trapped the sensor oxidation state with NEM (Figures 3 and 4B). The advantage here is that measurements are under fully physiological conditions at medium throughput. The disadvantage is that H_2_O_2_ levels are not monitored in real time and therefore rapid H_2_O_2_ dynamics may be missed. The decision of which setup to employ needs to be made depending on the specific biological question asked.

### Properties of the roGFP2-Tsa2ΔC_R_ sensor in Chlamydomonas cells

Overall, the kinetics of oxidation and reduction of roGFP2-Tsa2ΔC_R_ after the exogenous addition of H_2_O_2_ were very similar to those observed for this sensor in yeast (Morgan et al., 2016) and for the roGFP2-Orp1 sensor in Arabidopsis seedings (Ugalde et al., 2020). Consistent with previous observations in yeast, we observed that cytosolic roGFP2-Tsa2ΔC_R_ was more than 20% oxidized at steady state and that the oxidation of the untreated control sample decreased over time. In yeast, this decrease in sensor oxidation was shown to be caused by decreasing oxygen levels in the assay buffer (Morgan et al, 2016). Thus, our observations indicate that, as in yeast, roGFP2-Tsa2ΔC_R_ functions as an ultra-sensitive H_2_O_2_ sensor in Chlamydomonas that enables the monitoring of ‘basal’ cellular H_2_O_2_ levels. Although difficult to compare between organisms and experimental setups, we also note that whilst 0.1 mM of exogenously added H_2_O_2_ induced a strong roGFP2-Tsa2ΔC_R_ response in Chlamydomonas, 2 mM H_2_O_2_ was required to elicit a roGFP2-Orp1 response in Arabidopsis seedlings (Ugalde et al., 2020).

RoGFP2-Tsa2ΔC_R_ targeted to the ER was fully oxidized (Figure 2H). Full oxidation was also observed for an unfused roGFP2 targeted to the ER of tobacco leaf cells (Meyer et al., 2007) and for HyPer targeted to the ER in mammalian cells (Mehmeti et al., 2012). This is almost certainly due to the oxidative milieu in the ER lumen rather than to high ER lumenal H_2_O_2_ concentrations (Mehmeti et al., 2012). Specifically, the ER maintains an active disulfide generating machinery and a relatively oxidized glutathione pool (in the absence of glutaredoxins) that does not allow sensor reduction (Schwarzlander et al., 2016). Hence, in this compartment disulfide bond formation is largely favored over disulfide reduction.

Cytosolic roGFP2 alone was insensitive to exogenously added H_2_O_2_ and found to be constantly in a highly reduced state (Figure 2C). RoGFP2 has been shown to be in equilibrium with the 2GSH/GSSG redox couple, which is catalyzed by enzymatically active glutaredoxin (Gutscher et al., 2008; Marty et al., 2009; Morgan et al., 2013). This points to a large excess of GSH over GSSG in the Chlamydomonas cytosol, as is the case for the cytosol of a wide variety of other cell types (Schwarzlander et al., 2016).

HyPer has been reported to be susceptible to gene silencing beyond the cotyledon stage in Arabidopsis (Exposito-Rodriguez et al., 2013). Although silencing can be circumvented, for example by transient expression in *Nicotiana benthamiana* abaxial epidermal cells (Exposito-Rodriguez et al., 2017), this is not ideal. We found no silencing of our roGFP2-Tsa2ΔC_R_ constructs over a period of 1–2 years in Chlamydomonas. However, although we employed all ‘tricks’ for high-level transgene expression, which included using the strongest promoter currently available (*HSP70A-RBCS2*), a codon-optimized ORF containing the three *RBCS2* introns, the *RPL23* terminator, and the UVM4 expression strain (Schroda, 2019), sensor expression was only just sufficient to achieve a good signal-to-noise ratio in most compartments. When expression levels were reduced by about half, i.e. when the *PSAD* promoter was used instead of the *AR* promoter, or when the sensor was targeted to the thylakoid lumen, the signal became too noisy for reliable measurements. Hence, for future applications of this sensor, care must be taken that expression levels in the chosen Chlamydomonas strain are sufficiently high.

### H_2_O_2_ produced by PET in the stroma under physiological conditions readily diffuses into the cytosol but not to other subcellular compartments

Our observations demonstrated the suitability of roGFP2-Tsa2ΔC_R_ to monitor H_2_O_2_ levels under physiologically relevant conditions. For example, under high light intensities (1000 μmol photons m^−2^ s^−1^) we observed a rapid, PET-dependent increase in stromal H_2_O_2_ levels, together with a concomitant increase in cytosolic H_2_O_2_. Changes in H_2_O_2_ in the mitochondrial matrix were barely detectable and no change was observed in the nucleus. We also could not detect an increase in H_2_O_2_ levels in the nucleus, even when we increased the light intensity to 1500 μmol photons m^−2^ s^−1^ (Figures 3D, 3E). But we could detect H_2_O_2_ in the nucleus and mitochondria of cells supplemented with paraquat and/or metronidazole at low light intensities (Figure 3E). Thus, our findings partially corroborate earlier studies, which reported leakage of PET-dependent H_2_O_2_ from the chloroplast into other compartments (Mubarakshina et al., 2010; Caplan et al., 2015; Exposito-Rodriguez et al., 2017; Ugalde et al., 2020). However, our results also indicate that the H_2_O_2_ scavenging capacity of the cytosol is sufficiently high to quench the comparably low H_2_O_2_ concentrations originating from high light exposure, which effectively limits H_2_O_2_ diffusion into other subcellular compartments. In contrast, the cytosolic H_2_O_2_ scavenging capacity appears to become overwhelmed by the high H_2_O_2_ concentrations produced in the presence of metronidazole or paraquat (or exogenously added H_2_O_2_) and therefore H_2_O_2_ is not prevented from reaching other subcellular compartments. In summary, our results strongly support the conclusion that cellular H_2_O_2_ scavenging enzymes limit the intracellular diffusion of H_2_O_2_ thereby leading to the establishment of steep intracellular H_2_O_2_ concentration gradients. This interpretation would be in line with the observation that efficient transfer of H_2_O_2_ from chloroplasts to the nucleus in plant cells required that the two compartments are in close proximity. This is possible because of the mobility of chloroplasts in higher plants cells and appears unlikely for the architecture of Chlamydomonas cells with a single, large chloroplast.

### H_2_O_2_ is produced rapidly under heat stress and does not derive from PET

We found H_2_O_2_ levels to increase rapidly and transiently during heat stress (Figure 4). Rapid accumulation of H_2_O_2_ has been reported previously in tobacco seedlings, spinach leaves, or Arabidopsis and tobacco cell cultures when exposed to heat stress and therefore appears to be a conserved response in plant cells (Foyer et al., 1997; Vacca et al., 2004; Volkov et al., 2006; Gómez et al., 2008). In Chlamydomonas, H_2_O_2_ levels closely paralleled the temperature change in the culture and therefore H_2_O_2_ appears to be derived from a constitutive temperature-dependent cellular process (Figure 4A). As the increase in H_2_O_2_ levels was also observed in the presence of DCMU, it cannot derive from PET. However, there are many alternative production sites for H_2_O_2_ in plant cells that could be temperature controlled, such as the mitochondrial respiratory chain, limited substrate oxidases (with glycolate, xanthin, urate, sulfite, mono- and polyamines as substrates), type III peroxidases, or NADH oxidases in the plasma membrane (Cheeseman, 2007; Sun and Guo, 2016). Photorespiration is an unlikely source for H_2_O_2_ in Chlamydomonas, as it depends on input of electrons from PET and Chlamydomonas has a glycolate dehydrogenase that does not produce H_2_O_2_, rather than a glycolate oxidase (Kern et al., 2020).

Despite the rapid increase in H_2_O_2_ levels upon heat stress, Chlamydomonas cells induced the expression of scavenging enzymes such as superoxide dismutases, catalase, peroxiredoxins, or dehydroascorbate reductase only between 3 h and 24 h after the temperature shift (Mühlhaus et al., 2011; Hemme et al., 2014; Schroda et al., 2015). This indicates that the increased levels of H_2_O_2_ upon heat exposure are not detrimental to the cells during the first hours at elevated temperatures and might play a role in signaling.

In summary, we report on the generation of constructs enabling the expression of the ultra-sensitive H_2_O_2_ probe, roGFP2-Tsa2ΔC_R_, in five different subcellular compartments in Chlamydomonas. We show that these sensors respond readily to both exogenously added and endogenously produced H_2_O_2_ and reveal that, by following the response of sensors targeted to multiple subcellular compartments, the existence of intracellular H_2_O_2_ gradients can be inferred. We anticipate that the future application of these sensors will allow exciting new insights into subcellular H_2_O_2_ homeostasis and dynamics.

## Methods

### Strains and Culture Conditions

#### Chlamydomonas reinhardtii

UVM4 cells (Neupert et al., 2009) were grown in Tris-Acetate-Phosphate (TAP) medium (Kropat et al., 2011) on a rotatory shaker at a constant light intensity of ~40 μmol photons m^−2^ s^−1^. For high-light exposure, cells were grown to ~1×10^6^ cells/ml, transferred to an open 1-L beaker, placed on an orbital shaker, and exposed to 1000 or 1500 μmol photons m^−2^ s^−1^ provided by CF Grow® (CXB3590-X4). Transformation was done with the glass beads method (Kindle, 1990) as described previously (Hammel et al., 2020) with constructs linearized with NotI. Transformants were selected on 100 μg ml^−1^ spectinomycin.

### Cloning of sensor and signal peptide coding sequences

The roGFP2-Tsa2ΔC_R_ sequence (Morgan et al., 2016) was reverse translated using the most-preferred Chlamydomonas codons and equipped with the three Chlamydomonas *RBCS2* introns as suggested previously for foreign genes (Schroda, 2019), but using the native flanking sites of these introns (CAA/In1/GAT, ACG/In2/GCT, and TGC/In3/CTG). An AAG Lys codon was converted into the suboptimal AAA Lys codon to eliminate a GAAGAC BpiI recognition site. The sequence was flanked with BsaI cleavage sites such that upon BsaI digestion a 2132-bp fragment with AATG and TTCG overhangs is generated for the B3/4 position of level 0 parts according to the Modular Cloning (MoClo) syntax for plant genes (Weber et al., 2011; Patron et al., 2015) (Figure 1A). Synthesis and cloning into the pUC57 vector was done by GeneCust (Luxembourg), giving rise to pMBS419. The latter was used as template for PCR with primers 5’-TT<u>GAAGAC</u>ATAATGGCCAGCGAGTTCAGC-3’ and 5’-TT<u>GAAGAC</u>TCCGAACCCTTGTACAGCTCGTCCATGCC-3’ to amplify a 898-bp fragment containing only the roGFP2 coding sequence with the first *RBCS2* intron (BpiI recognition sites are underlined). The PCR product was combined with destination vector pAGM1287 (Weber et al., 2011), digested with BpiI and assembled by ligation into level 0 construct pMBS467. The level 0 part encoding the HSP70B chloroplast transit peptide was generated via PCR on plasmid pMBS197 using primers 5’-TT<u>GAAGAC</u>AACCATGCCGGTTCAGCAGATGAC-3’ and 5’-TT<u>GAAGAC</u>AACATTGCCTGAAGGAACAATTCAAATG-3’. This plasmid harbors the coding sequence for the HSP70B transit peptide with an intron, in which a BsaI recognition site had been eliminated. The resulting 251-bp product and plasmid pAGM1276 (Weber et al., 2011) were digested with BpiI and ligated to yield pMBS639. The level 0 part for the chloroplast transit peptide of CDJ1 was produced accordingly by PCR using primers 5’-TT<u>GAAGAC</u>AACCATGCTCGCAAACCTTCGTAG-3’ and 5’-TT<u>GAAGAC</u>AACATtCCGTCGGCCCGCACAACCAC-3’ on genomic DNA, generating a 166-pb product and giving rise to pMBS640. The same procedure was used to produce level 0 parts for bipartite transit peptides for stroma and thylakoid lumen of PSAN and PSBO, using primers 5’-AA<u>GAAGAC</u>CACCATGGCCATCTCTGCTCGCTC-3’ and 5’-TT<u>GAAGAC</u>GGCATTCCGGCGTTGGCGACGGGGGC-3’, and primers 5’-TT<u>GAAGAC</u>AACCATGGCCCTCCGCGCTGCCCA-3’ and 5’-TT<u>GAAGAC</u>AACATTGCgAGGGCGTTGGCCGACTGCGA-3’ to generate PCR products of 508 bp and 470 bp, giving rise to pMBS298 (PSAN) and pMBS641 (PSBO), respectively. All PCRs were done with Q5 High-Fidelity Polymerase (NEB) following the manufacturer’s instructions. Correct cloning was verified by Sanger sequencing. These newly constructed level 0 parts were then complemented with level 0 parts (pCM) from the Chlamydomonas MoClo toolkit (Crozet et al., 2018) to fill the respective positions in level 1 modules as follows: A1-B1 – pCM0-015 (*HSP70A-RBCS2* promoter + 5’UTR) or pCM0-010 (*PSAD* promoter + 5’UTR); A1-B2 – pCM0-020 (*HSP70A-RBCS2* promoter + 5’UTR) or pCM0-016 (*PSAD* promoter + 5’UTR); B2 – pCM0-053 (USPA chloroplast transit peptide), pMBS639 (HSP70B chloroplast transit peptide), pMBS640 (CDJ1 chloroplast transit peptide), pMBS298 (PSAN thylakoid lumen targeting peptide), pMBS641 (PSBO thylakoid lumen targeting peptide), pCM0-057 (HSP70C mitochondrial transit peptide), or pCM0-056 (BIP ER targeting signal); B3/4 – pMBS419 (roGFP2-Tsa2ΔC_R_) or pMBS467 (roGFP2); B5 – pCM0-100 (3xHA), pCM0-101 (MultiStop), pCM-111 (BIP ER retention sequence), or pCM-109 (SV40 nuclear localization signal); B6 – pCM0-119 (*RPL23* 3’UTR). The *HSP70A-RBCS2* fusion promoter used here contains −467 bp of *HSP70A* sequences upstream from the start codon in optimal spacing with respect to the *RBCS2* promoter (Lodha et al., 2008; Strenkert et al., 2013). The respective level 0 parts and destination vector pICH47742 (Weber et al., 2011) were combined with BsaI and T4 DNA ligase and directionally assembled into the eleven level 1 modules shown in Figure 1B. The level 1 modules were then combined with pCM1-01 (level 1 module with the *aadA* gene conferring resistance to spectinomycin flanked by the *PSAD* promoter and terminator) from the *Chlamydomonas* MoClo kit, with plasmid pICH41744 containing the proper end-linker, and with destination vector pAGM4673 (Weber et al., 2011), digested with BpiI, and ligated to yield the eleven level 2 devices displayed in Figure 1B. All newly generated level 0 and level 2 plasmids can be ordered from the Chlamydomonas Research Center (https://www.chlamycollection.org/).

### Protein analysis by SDS-PAGE

Protein extractions, SDS–PAGE, semi-dry blotting and immunodetections were carried out as described previously (Hammel et al., 2020). Antibodies used for immunodetection were mouse anti-HA (Sigma H9658, 1:10,000) and rabbit anti-GFP (Roche, Cat. No. 11814460001). Densitometric band quantifications after immunodetections were done with the FUSIONCapt Advance program (PEQLAB).

### RoGFP2 fluorescence recordings

RoGFP2 fluorescence was recorded using a CLARIOstar fluorescence plate reader (BMG-Labtech) as described previously for yeast (Morgan et al., 2016) and adopted to Chlamydomonas as follows: Chlamydomonas cells were grown at constant light to a density of ~3×10^6^ cells/ml. 1×10^7^ cells were used per well and were harvested by centrifugation at 3800 rpm for 2 min at 25°C. Cells were resuspended in 100 mM MES-Tris buffer (pH 7.0) (5×10^7^ cells/ml) and 200 μl cell suspension were transferred per well of a black, flat-bottomed 96-well imaging plate (BD Falcon 353219). For calibration and later data processing, control samples of fully oxidized and fully reduced sensors were prepared by adding N,N,N’,N’-tetramethylazodicarboxamide (diamide) at a final concentration of 20 mM and DTT at a final concentration of 100 mM to cells in two different wells. Further wells to record roGFP2 fluorescence at steady state, at increasing H_2_O_2_ concentrations, or after different treatments were prepared. Before starting the measurement, the 96-well imaging plate was centrifuged at 30 g for 5 min at 25°C. Chlamydomonas UVM4 recipient cells were always included as negative control. The measurement mode of the plate reader was based on the “Fluorescence Intensity” coupled with the “Bottom Optic” option, with a positioning delay of 0.2 sec, 40 flashes per well and the desired number of measurement cycles (about 200 cycles) and time per cycle (about 20 sec). The gain adjustment was set to 80% maximum and an automatic gain adjustment for each wavelength was performed immediately before the measurement. RoGFP2 exhibits two excitation maxima at 400 nm and 475–490 nm when fluorescence emission is monitored at 510 nm. Therefore, it was ensured that the number of multichromatics was set to 2 and that the correct excitation (410 and 480 nm) and emission (510 nm) filters were selected. The measurement was started and increasing H_2_O_2_ solutions were added after 10 cycles. After the fluorescence measurement was done, MARS Data Analysis software was used to subtract the auto-fluorescence of WT cells. The “blank corrected” values were taken to calculate the degree of sensor oxidation (OxD) using the following equation:

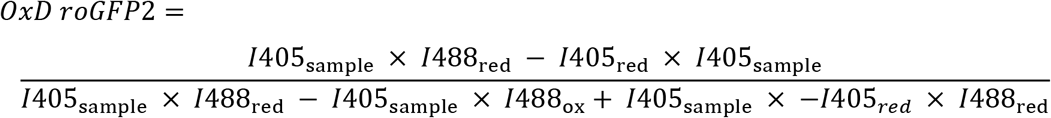

where *I* is the fluorescence intensity at 510 nm after excitation at either 405 nm or 488 nm. The subscripts ‘ox’, ‘red’ and ‘sample’ indicate the fluorescence intensity measured for the fully oxidized and fully reduced controls and the sample, respectively.

### NEM-based alkylation for measuring H_2_O_2_ levels in cell cultures

The cell number of Chlamydomonas cells grown to 1-3×10^6^ cells/ml was determined and the volume to harvest 2.5×10^7^ cells was calculated. For rapid ‘trapping’ of probe oxidation, 50 ml Falcon tubes were preloaded with one fourth of the final volume with 100 mM MES-Tris (pH 7.0) buffer containing 40 mM NEM. Chlamydomonas cells were harvested in prefilled Falcon tubes by centrifugation at 3800 rpm for 3 min at room temperature and subsequently resuspended in 500 μl 100 mM MES-Tris (pH 7.0) buffer containing 40 mM NEM to a final concentration of 5×10^7^ cells/ml. 200 μl (10^7^ cells per well) of suspension were transferred to a 96-well plate. For calculating the OxD, fully oxidized (diamide) and reduced (DTT) samples were added and probe oxidation was measured following the protocol described above.

### Continuous monitoring of real-time changes in H_2_O_2_ during heat stress via connected tubing and a peristaltic pump

Chlamydomonas cells (~5×10^6^ cells/ml) were cultivated under agitation at a constant light intensity of ~40 μmol photons m^−2^ s^−1^ and continuously circulated to a quartz cuvette inserted into a spectrofluorometer (Jascon FP-8300) via connected tubing and a peristaltic pump (Watson Marlow 101U, flow rate: 1 ml/s). After 20 min, cells were exposed to 40°C for 30 min in a water bath, and shifted back to 23°C for 30 min. The ratio of the fluorescent light intensities emitted after excitation at 405 and 488 nm was monitored every 30 s. The measurement mode of the spectrofluorometer was based on the “Fixed wavelength measurement” with an excitation bandwidth of 5 nm and an emission bandwidth of 10 nm.

### Confocal microscopy

All images were acquired using a Zeiss LSM880 AxioObserver confocal laser scanning microscope equipped with a Zeiss C-Apochromat 40x/1,2 W AutoCorr M27 water-immersion objective. Fluorescent signals of green fluorescent protein (GFP) (excitation/emission 488 nm/491-589 nm) and chlorophyll autofluorescence (excitation/emission 633 nm/647-721 nm) were processed using the Zeiss software ZEN 2.3 or ImageJ.

---

**Supplemental Figure 1.** Screening of transformants expressing the roGFP2 sensor by immunoblotting.

**Supplemental Figure 2.** Analysis of fluorescence properties of the roGFP2-Tsa2ΔCR sensor at different temperatures.

## Acknowledgements

This work was supported by the Deutsche Forschungsgemeinschaft [SFB/TRR175, project C02] and the Profilbereich BioComp. We would like to thank Nicole Frankenberg-Dinkel for providing access to her confocal microscope and her spectrofluorometer and Johannes Herrmann for providing access to his fluorescence plate reader.

## Author Contributions

J.N. generated all constructs and performed all experiments. D.S recorded the confocal microscopy images. J.O. helped with setting up the fluorescence recordings. M.S. and B.M. conceived and supervised the project and wrote the article with contributions from all authors.

